# A new spin on chemotaxonomy using non-proteogenic amino acids as a test case

**DOI:** 10.1101/2024.09.28.615597

**Authors:** Makenzie Gibson, William Thives Santos, Alan R. Oyler, Lucas Busta, Craig A. Schenck

**Author notes:** Equal contribution (Makenzie Gibson, William Thives Santos).

## Abstract

**Premise:** Specialized metabolites serve various roles for plants and humans. Unlike core metabolites, specialized metabolites are restricted to certain lineages. Thus, in addition to their ecological functions, specialized metabolites can serve as diagnostic markers of plant lineages.

**Methods:** We investigate the phylogenetic distribution of plant metabolites using non-proteogenic amino acids (NPAA). Species-NPAA associations for eight NPAAs were identified from the existing literature and placed within a phylogenetic context using R packages and interactive tree of life. To confirm and extend the literature-based NPAA distribution we selected azetidine-2-carboxylic acid (Aze) and screened over 70 diverse plants using GC-MS.

**Results:** Literature searches identified > 900 NPAA-relevant articles, which were manually inspected to identify 560 species-NPAA associations. NPAAs were mapped at the order and genus level, revealing that some NPAAs are restricted to single orders, whereas others are present across divergent taxa. The distribution of Aze across plants suggests a convergent evolutionary history.

**Discussion:** The reliance on chemotaxonomy has decreased over the years. Yet, there is still value in placing metabolites within a phylogenetic context to understand the evolutionary processes of plant chemical diversification. This approach can be applied to metabolites present in any organism and compared at a range of taxonomic levels.

## INTRODUCTION

Plants are remarkable for their chemical diversity. These metabolites enable plants to interact with their environment and provide various services to humans including medicines (Li and Weng, 2017; Weng et al., 2021). Roughly one third of medicines are derived from plants and much of the global population relies directly on plants as medicinal sources (McChesney et al., 2007; Newman and Cragg, 2016). As a group, plants make approximately 200,000 distinct metabolites and other estimates are as high as 1 million (Dixon and Strack, 2003; Rai et al., 2017; Alseekh and Fernie, 2018). Yet, much of the plant chemical diversity remains undiscovered. This underscores the relatively untapped potential of plants as a source for new chemistries particularly useful in human health and disease treatment.

In addition to core metabolism, which encompasses essential pathways such as membrane, nucleotide, and protein biosynthesis, plants also produce specialized metabolites. While specialized metabolites are not strictly necessary, they enable interactions with the environment (Erb and Kliebenstein, 2020). Another hallmark of plant specialized metabolites is their restricted distribution across plants (Moghe and Last, 2015; Schenck and Last, 2020). Unlike core metabolites that are ubiquitous, some specialized metabolites are produced in a narrow range of plants or a single species (e.g. morphine; Beaudoin and Facchini, 2014), produced by related species within the same family (e.g. acylsugars; Vendemiatti et al., 2024), or accumulated in unrelated plants (e.g. caffeine; Huang et al., 2016). Thus, metabolites can serve as diagnostic traits in plant taxonomy.

Chemotaxonomy is the field of merging chemistry and systematics (Alston et al., 1963; Gibbs, 1974; Reynolds, 2007). Early literature focused on building taxonomic tress based on abundant plant metabolites (Bate-Smith, 1962). While useful for some metabolites, it was impractical given the numerous instances of convergent evolution of plant metabolic pathways of which we are now aware (Pichersky and Lewinsohn, 2011). Although still useful, it is now more prescient to map metabolites onto existing high-quality plant phylogenies based on sequence data, which are both abundant and of high quality (Qian and Jin, 2016; Leebens-Mack et al., 2019; Zuntini et al., 2024). Mapping metabolic traits into existing phylogenies provides an indication of how metabolic pathways emerge, enables hypotheses about convergent evolution, and provides new directions for the study of plant lineages and their metabolism.

Non-proteogenic amino acids (NPAAs) are amino acids by definition, with amino and carboxyl groups bound to an alpha carbon, but are not typically incorporated during protein biosynthesis, unlike the canonical 20 proteogenic amino acids. Some NPAAs are widely distributed across plants and act as intermediates in core metabolism such as S-adenosylmethionine and ornithine (Bell, 2003; Huang et al., 2011; Vranova et al., 2011). However, most NPAAs are restricted to certain lineages and thought to serve defensive roles (Huang et al., 2011; Vranova et al., 2011). Many NPAAs are structural analogs of proteogenic amino acids and can be misincorporated during protein biosynthesis, though mechanisms of action are not well studied (Rodriguez-Mias et al., 2022; Thives Santos et al., 2024). Greater understanding of the distribution of NPAAs can inform studies on biosynthetic pathways and biochemical processes related to their mechanisms of action.

Here, we put a new spin on chemotaxonomy using NPAAs as a test class of metabolites. We leverage the abundant literature that exists for detection of NPAAs in plants and high-quality nucleotide-based phylogenies. We focus on eight NPAAs that have been detected across multiple plants including, azetidine-2-carboxylic acid (Aze), canaline, canavanine, djenkolic acid, 5-hydroxytryptophan, indospicine, meta-tyrosine, and mimosine. We mined the literature using text mining and manual curation to establish a near-comprehensive list of species-NPAA associations, which were then mapped onto existing plant phylogenies at varying taxonomic scales using R packages and interactive tree of life (iTOL). To confirm and extend the species-NPAA associations from the literature, we extracted and detected Aze from more than 70 diverse plants. Our results were consistent with the literature and suggest that Aze likely evolved through convergent evolution in divergent taxa. This study provides a template that can be applied to any (plant) metabolite, which can be used to understand the emergence of metabolic pathways.

## METHODS

### NPAA-plant associations in existing literature

To understand the phylogenetic distribution of NPAAs we mined the literature to identify species-NPAA associations. We focused on eight NPAAs that have been detected across multiple plants including azetidine-2-carboxylic acid, canaline, canavanine, djenkolic acid, 5-hydroxytryptophan, indospicine, meta-tyrosine, and mimosine because of their occurrence throughout the literature and their restricted phylogenetic distribution (Bell 1976; Bell 2003, Fowden 1963, Huang et al., 2011). A comprehensive list of 900 publications were identified using text mining searches that contained potential species-NPAA associations. These were manually curated to remove literature that were inadvertently captured in text mining searches and to remove duplicates, correct for usage of common names and genus species name changes to identify a confident list of plant-metabolite associations (Supplementary Table 1).

### Phylogenetic mapping of NPAAs

Plant-metabolite associations were mapped onto an existing plant megaphylogeny (Qian et al., 2016). We pruned the megaphylogeny to show only certain lineages using the “buildTree” function available at https://github.com/thebustalab/phylochemistry, then visualized trees using the R package ‘ggtree’ (Yu et al., 2017). Metabolite associations were plotted alongside phylogenies using ggplot2 or integrative tree of life (iTOL) (Wickham, 2009; Letunic and Bork, 2024). For the species tree, a total of 789 species are represented on the tree, 395 from species with NPAA associations and 394 randomly selected species. All code used in our analyses is available at https://github.com/thebustalab/npaa_distribution.

### Aze detection from diverse plants

To confirm and extend the literature-based approach, Aze was extracted, detected and analyzed from various plant lineages. Plant material was obtained through multiple sources; fresh leaf tissues from The Missouri Botanical Garden in St. Louis, MO, fresh leaf tissues from the University of Missouri Botanical Garden, Columbia, MO, or from plant tissues grown from seeds obtained from the USDA germplasm resource. In most cases fresh leaf material was collected and flash frozen in liquid nitrogen, lyophilized for at least 48 hours until completely dry and stored until analysis. Supplementary Table 2 contains a list of all species, tissues and source of the plant material. Glass beads were placed in a 15 or 50mL conical tube together with dry plant material and ground to a fine powered using a bead mill (Spex Geno/grinder). Seeds were used in some cases, legume lineages, because germination failed. Seeds were ground to a fine powder using a Perten Labmill 3310. Between 10-20 mg of finely ground plant material was used for extraction using 700 μL of water. Samples were incubated for 1 hour at 50 °C with vortexing every 15 minutes. 700 μL of chloroform was added to the sample and incubated for 1 hour at 50 °C with vortexing every 15 minutes. Extracts were spun at 3000 x g for 30 minutes at 4 °C and 650 μL of the top layer was added to a glass vial and dried to completeness using a speedvac (Labconco). The dried extract was resuspended in 50 μL of a pyridine solution containing 15mg/mL of methoxyamine HCl and incubated for 1 hour at 50 °C, then 50 μL of *N*-methyl-*N*-(*tert*-butyldimethylsilyl) trifluoroacetamide + 1% tert-butyldimetheylchlorosilane (MTBSTFA + 1 % t-BDMCS, Supelco) was added and incubated for 1 hour at 50 °C. Derivatized samples were injected onto an Agilent 5977C GC-MS with authentic Aze standard (Sigma). 1uL of sample was injected with a 5:1 split ratio onto a 60m DB-5ms column (Agilent). The initial oven temperature was 120 °C with an oven ramp of 6 °C/min until 300 °C and held for 8 minutes. The inlet valve temperature was maintained at 280 °C. The MS was operated in full scan mode from m/z 50-650. A limit of detection for Aze was determined through injection of a concentration gradient of Aze from 0-1 mM, yielding a limit of detection of 57.2 μM. RESULTS:

### Identification of NPAA-species associations

To determine the phylogenetic distribution of NPAAs, we first mined the literature to identify species-metabolite associations. NPAAs are a good class of metabolites to use as a framework for phylochemical mapping because there has been substantial interest in these compounds starting around the 1950s, and there is a large base of literature that contains NPAA detections across phylogenetically diverse plants (Grobbelaar et al., 1955; Fowden and Steward, 1957; Bell, 1976). We focused on eight NPAAs including azetidine-2-carboxylic acid (Aze), canaline, canavanine, djenkolic acid, 5-hydroxytryptophan, indospicine, meta-tyrosine, and mimosine (Fig. 1a) because of their occurrence throughout the literature and their restricted phylogenetic distribution. These structurally diverse NPAAs also have distinct mechanisms of action, some mimic proteogenic amino acids and are misincorporated during protein biosynthesis (Aze, m-Tyr, canavanine (Bertin et al., 2007; Zer et al., 2020; Thives Santos et al., 2024), some react with endogenous metabolites and cause adverse effects (canaline; Rosenthal 1997), while others remain relatively unknown (indospicine, djenkolic acid). Literature searches identified more than 900 scientific articles with mentions of NPAAs. Manual curation of these literature reports identified 560 species-NPAA associations (Supplementary Table 1). A list of all sources used to identify species-metabolite associations can be found in Supplementary Table 1. Canavanine was the most frequently detected NPAA in the literature with 433 species associations and canaline was detected in the fewest with 3 associations identified (Supplementary Table 1). This near-comprehensive list of select NPAA species associations could then be used as the framework for phylogenetic mapping of the metabolic traits.

**Fig. 1.**
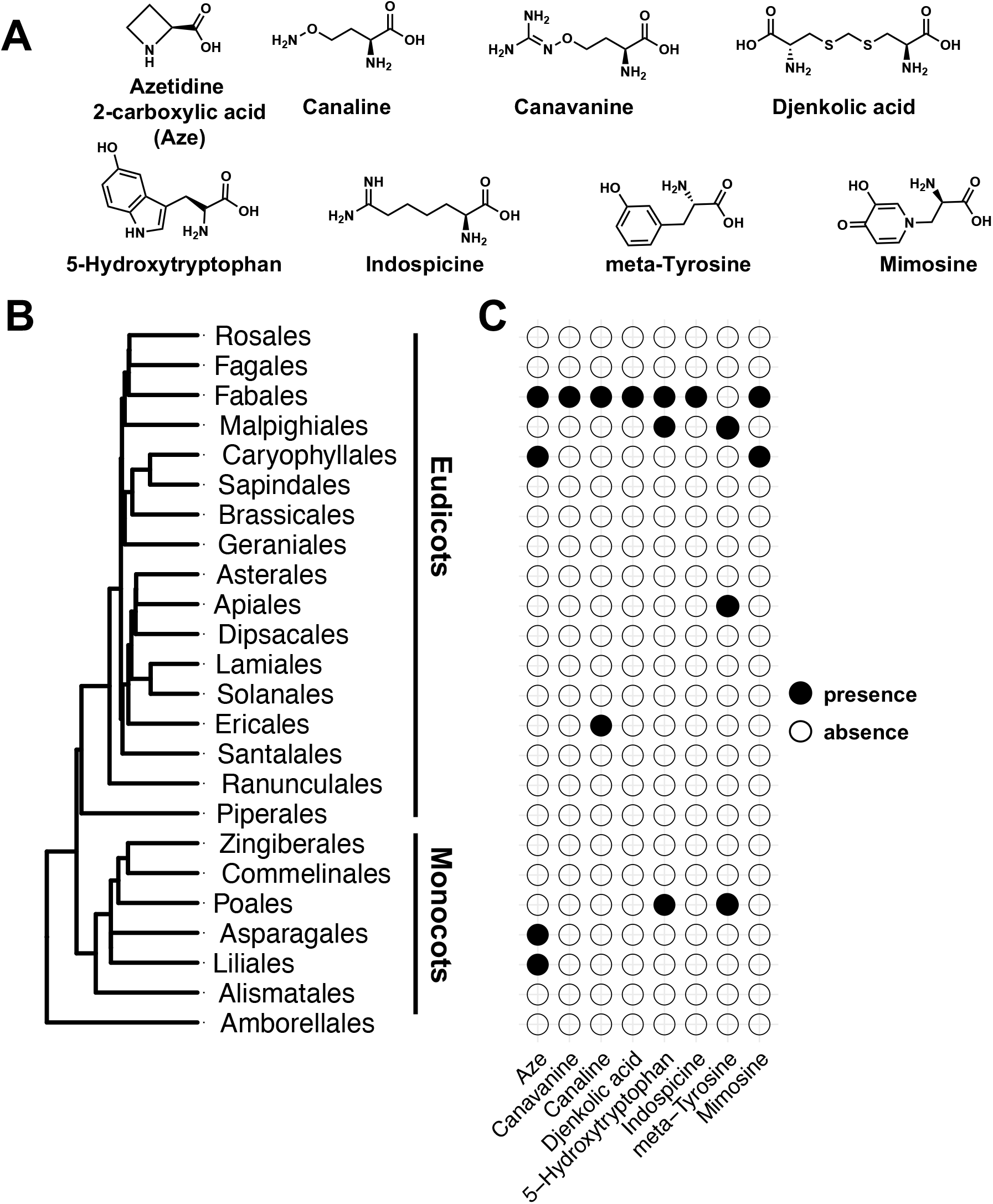
Order level distribution of NPAAs across plants. **(A)** Structures of the eight NPAAs that were a focus of this study. **(B)** Order level phylogeny of flowering plants. 23 orders were selected that represented the diversity of flowering plants and a sister lineage (Amborellales). **(C)** Phylogenetic mapping of NPAA-species associations. Orders with species that contain a particular NPAA-species association in the literature are represented by a filled circle.

### NPAA distribution at the order level

To understand the distribution of the eight NPAAs at a global scale, order level phylogenies were constructed (Qian and Jin, 2016). In total 23 orders were selected representing the diversity of flowering plants and used as a basis for mapping NPAAs using ggtree (Fig. 1B; Yu et al., 2017). NPAA-species associations were not identified for most of the orders on the phylogeny, but provide a sense for the distribution of NPAAs across plants at a global scale. Aze, canaline, canavanine, djenkolic acid, 5-hydroxytryptophan, indospicine, meta-tyrosine, and mimosine were then plotted into this phylogeny using ggplot2 (Wickham, 2009). In general, the NPAAs show a narrow distribution and are only found in a few orders (Fig. 1C). The Fabaceae is well known for the accumulation of diverse NPAAs (Bell et al., 2008), and consistently, 7 of the 8 NPAAs were found to associate with Fabales (Fig. 1C). Some NPAAs were reported in only a single order and show a very narrow distribution, such as canaline and djenkolic acid (Fig. 1C). Despite canavanine being identified in over 400 species, all these were within the Fabales order (Fig. 1C). Whereas, the other NPAAs were found in a few unrelated orders, such as mimosine in the Fabales and Caryophyllales and 5-hydroxytryptophan in the Fabales, Malpighiales, and Poales (Fig. 1C). The NPAAs that are reported in distinct orders could be examples of metabolites that have evolved through convergent evolution, but finer mapping and a deeper survey of species could provide more support for this hypothesis. It should be noted that lack of a literature report of a NPAA does not indicate absence of the metabolite in that plant. Thus, although it appears as though NPAAs show limited distribution, this reflects sampling bias and lack of data. Mapping of NPAAs at the order level gives a global view of the distribution, but finer scale mapping at the genus or species taxonomic levels could provide greater resolution and insight into the evolutionary trajectory of the underlying metabolic pathways.

### Species level NPAA distribution

To gain a finer resolution of the distribution of the selected NPAAs across plants, we mapped distributions into a species tree. Here we used all species that had at least one NPAA detected (560) and 500 randomly selected species to enrich the phylogenetic analysis. Our species tree consists of 789 species because not all the species with NPAA associations or the randomly selected species were present in the megaphylogeny we used as a source to prepare our species-level phylogenetic tree (Fig. 2). Species names were removed for interpretability, however a complete tree with species labels is present in Supplementary Figure 1. Species with a box indicate that a particular NPAA has been reported in the literature for that species. Species without boxes indicates lack of data and not necessarily that the plant does not accumulate the NPAA.

**Fig. 2.**
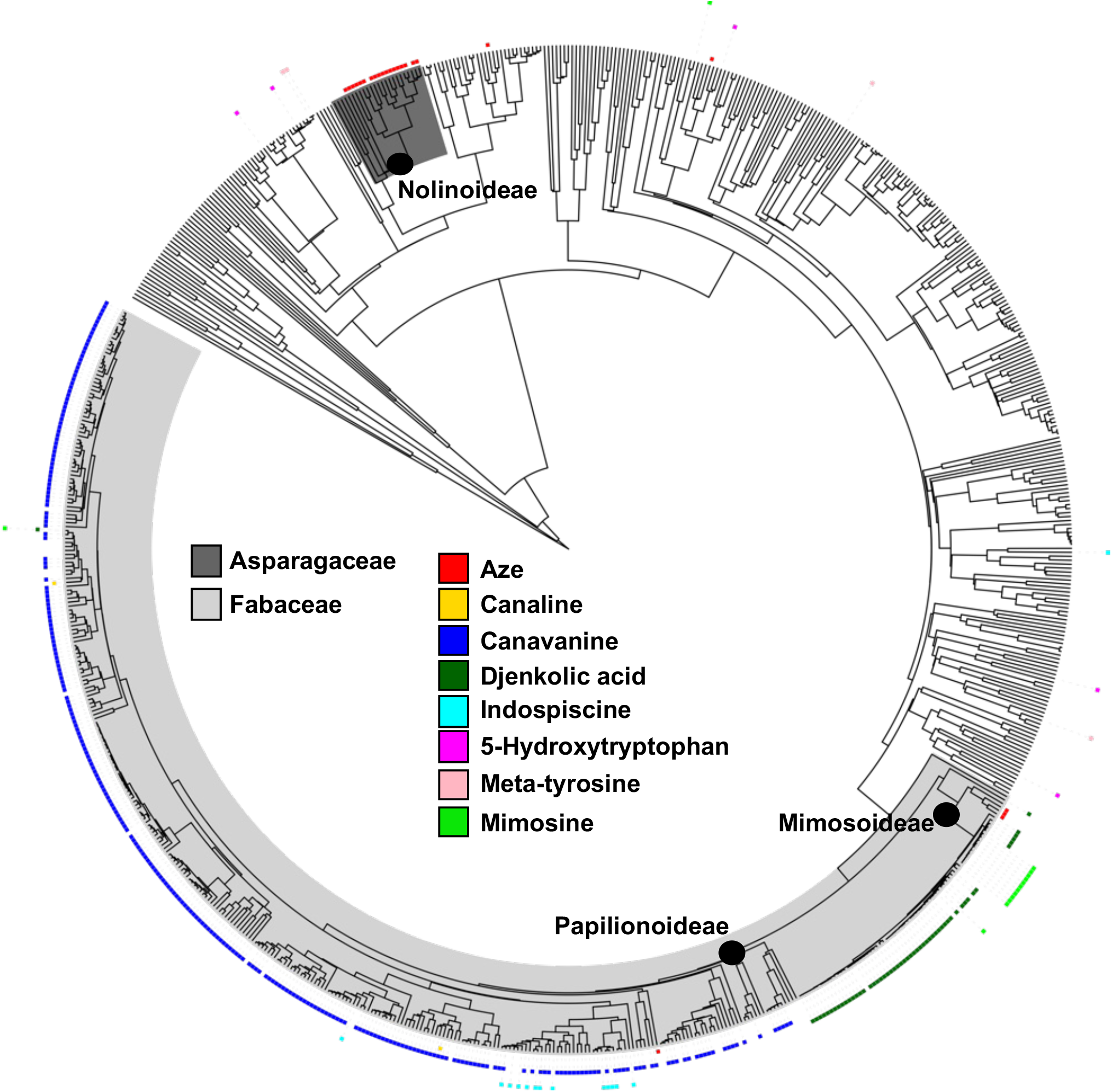
Species level distribution of NPAAs across plants. A total of 560 species-NPAA associations were identified in the literature. A species level phylogeny was built using these 560 species and supplemented with 500 random species. The final tree contained 789 species. The 8 NPAAs were mapped onto this tree. Plant names are removed for clarity. A full tree can be found in (Supplementary Fig. 1). Individual NPAAs are plotted onto the corresponding species with a filled box. Major plant clades are labeled on the tree including families and subfamilies mentioned in the main text.

Canavanine has been detected in the most species (433 canavanine-species associations; Supplementary Table 1), but all are restricted to the Fabales order (Fig. 1). Within the Fabales, canavanine appears to be restricted to the Papilionoideae subfamily, which contains species such as alfalfa and common bean (Fig. 2). Based on this phylogeny, it is likely that canavanine evolved in the common ancestor that has given rise to the modern day Papilionoideae subfamily Gibson and Thives Santos et al., *Non-proteogenic amino acid distribution across plants* and likely been retained in many, if not all lineages. We found 56 djenkolic acid-species associations (Supplementary Table 1). Djenkolic acid has only been detected in legume species (Fig. 1C) mostly within the *Acacia* genus with two exceptions, however all within the Mimosoideae subfamily (Fig. 2). Djenkolic acid likely emerged in the lineage that has given rise to *Acacia* and been retained in most, if not all, modern-day *Acacia* lineages (Fig. 2). Aze was identified in 28 species in the literature and in four distinct orders (Fig. 1; Supplementary Table 1), and within those orders is restricted in its distribution to a few closely related legumes in the Mimosoideae subfamily and species within the Nolinoideae subfamily of the Asparagales order (Fig. 2).

The other NPAAs have been reported in much fewer species. Mapping their distribution on a species tree highlights the need for more sampling prior to making inferences about how these pathways may have evolved. Mimosine was identified in 14 lineages, and all but one were within the legumes (Fig. 2). All these associations were found within the *Leucaena, Acacia*, and *Mimosa* genera (Fig. 2) showing a limited distribution. Indospicine was identified in 14 species with all but one report within the Fabales. Within Fabales indospicine is restricted to the *Indigofera* genus (Fig. 2). 5-hydroxytrptophan was detected in 7 species (Fig. 2). Limited sampling limits further interpretation within a species context, however 5-hydroxytrptophan is widely distributed and present in three orders (Fig. 1C). Meta-tyrosine had 5 associations across 3 orders and the only NPAA that was investigated that is not present within Fabales (Fig. 2). Canaline was detected in 3 plants and all but one were restricted to the legumes (Fig. 2). Only a handful of species were reported to contain multiple NPAAs, for example, both mimosine and djenkolic acid have been reported in *Mimosa pudica*, and canavanine and indospicine were both found within *Indigofera suffruticosa* (Fig. 2, Supplementary Table 1). This reflects the lack of a comprehensive dataset representing NPAA detection across a wide range of species, as most plants produce a suite of defense metabolites to target generalist and specialist predators (Endara et al., 2023), it is likely that plants accumulate multiple types of NPAAs.

### Validation of the literature and extension of NPAA-plant associations

There are limitations to our large-scale phylochemical mapping approach, such as inadvertently assigning NPAA-species associations that do not exist because of discrepancies in names, name changes or use of common names and there may be species with known NPAA accumulation that we have missed in our approach. To validate our literature-based NPAA phylochemical mapping approach, we chose Aze to determine definitive species-Aze associations by metabolite extraction and detection using GC-MS. Aze was first detected in *Convallaria majalis* (lily of the Valley (Fowden, 1955), and subsequently in additional plants within the Asparagales and Fabales orders (Fowden and Steward, 1957; Sung and Fowden, 1969). We also used this opportunity to further refine Aze distribution by screening plants closely related to Aze accumulators. In total we collected tissue from 78 species. Plants collected from botanical gardens were identified with family, genus, and species names and GPS coordinates to verify accurate collection, plants were also sourced from the USDA-ARS Germplasm Resource Information Network (GRIN) and locally (Supplementary Table 2). From the literature, Aze was detected 28 times, these associations were plotted into a genus tree (Fig. 3). The size of the circle indicates the number of species and reports of Aze detection within each genus. Aze was mainly detected within Asparagales and Fabales (Fig. 3), with reports of Aze also being detected from table beet within the genus *Beta* (Fig. 3).

**Fig. 3.**
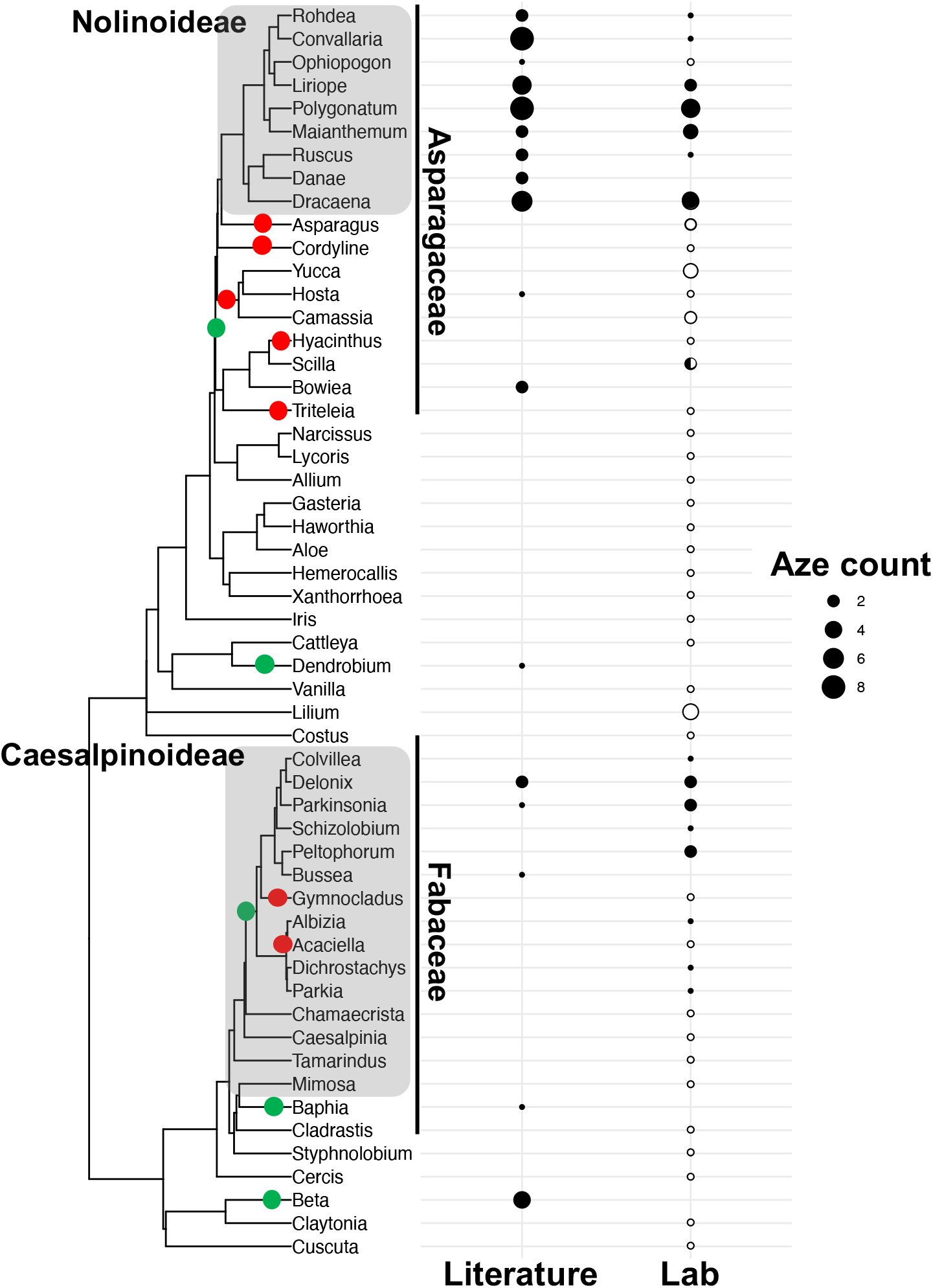
Confirmation and extension of literature-based Aze distribution. Aze was mapped onto a genus level phylogeny consisting mostly of plants within the Fabales and Asparagales orders (Literature). For literature data, the size of the circle indicates the number of species and literature reports within each genus, with larger circles indicating more Aze-species associations. 78 species were collected and analyzed for Aze using GC-MS (Lab). The size of the circle indicates number of species within each genus that were analyzed. Filled circles indicate presence of Aze and unfilled indicate absence. Circles with partial fill represent genera with some species that accumulate Aze and others where Aze was not detectable. Major plant clades are labeled next to the phylogeny. Green circles represent potential origins of Aze biosynthesis, whereas red circles represent potential loss events of Aze biosynthesis based on the literature and lab Aze distribution.

To isolate and detect Aze from diverse plants, we performed metabolite extractions primarily from leaf tissue, however other tissues were used as indicated in Supplementary Table 2, followed by derivatization and detection using GC-MS (Fig. 4). Methods were developed using authentic Aze standards and used to compare to plant extracts (Fig. 4). Our Aze analyses mostly confirmed what has been reported in the literature (Fig. 3). We also extend and refine the known distribution of Aze. For our analysis, filled circles indicate genera where Aze was detected, the size of the circle indicates the number of species within that genus that we detected Aze, e.g., a circle size 4 means that we detected Aze from 4 different species within that genus. Empty circles indicate genera that we tested, but did not detect Aze, and circles that are not entirely filled indicate genera that contained some species that accumulated Aze and others that did not, with the portion of the filling representative of the percent of species that accumulate Aze (Fig. 3). Our analysis shows that Aze is narrowly distributed in both the Fabaceae and Asparagaceae (Fig. 3). Initially we hypothesized that Aze was only found within the Nolinoideae subfamily because a sister genus, *Asparagus*, does not accumulate Aze in two species tested (Fig. 3). However, Aze was detected in two genera (*Bowiea* and *Scilla*), which were all outside the Nolinoideae subfamily, but within Asparagaceae (Fig. 3). These represent genera where wider sampling is needed to confirm Aze distribution. These data suggest that Aze biosynthesis likely emerged at the base of the Asparagaceae but has been lost in some lineages (Fig. 3 green and red dots, respectively). There is only one report of Aze outside of Asparagaceae, but within monocots within the *Dendrobium* genus (Fig. 3). We were unable to collect tissue for this genus, however we were able to collect tissue from closely related genera, and did not detect Aze in any of these lineages (Fig. 3). Aze detection in *Dendrodium* could represent an interesting case of independent evolution of Aze, or that this represents a false positive. There are only a few instances of discrepancies in the literature in our Aze analysis. Aze-species associations were identified for *Hosta* and *Ophiopogon*, however our analysis of a single *Hosta* and *Ophiopogon* species were unable to detect Aze (Fig. 3).

**Fig. 4.**
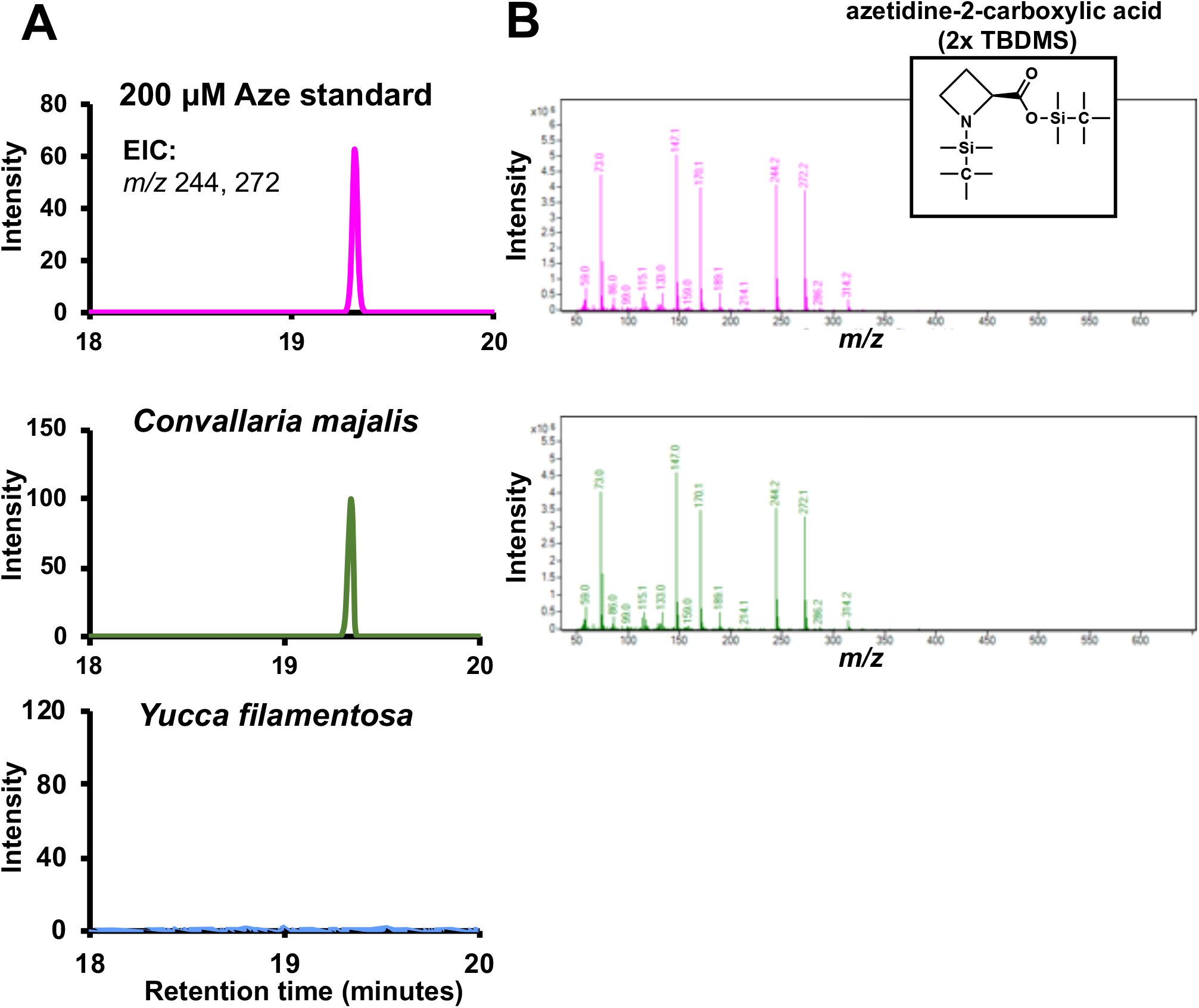
GC-MS detection of Aze from plant extracts. Metabolite extractions were performed from lyophilized plant tissue. Dried extracts were derivatized using MTBSFTA and analyzed using GC-MS. Authentic Aze standards were compared with plant extracts. **(A)** Extracted ion chromatograms for two ions (244 and 272) that are unique to Aze. Chromatograms show that the authentic Aze standard (pink) and extracts from *Convallaria majalis* (green) have a peak with the same retention time and **(B)** the same mass spectra. *Yucca filamentosa* (blue) represents a species where Aze was not detected. Inset shows derivatized Aze structure.

Aze detection in legumes was narrowly reported in the literature (Fig. 3). We extended this analysis and find that Aze is distributed more widely than initially thought. Aze is mainly found within the Caesalpinioideae subfamily, with one literature report outside but within Fabaceae (Fig. 3). Although we were unable to collect and screen tissue for a *Baphia* species, we did screen closely related legumes, and did not detect Aze in those lineages (Fig. 3). Aze distribution in Fabales suggests that biosynthesis likely emerged in the common ancestor shared by the Caesalpinioideae subfamily and there were a few loss events, for example in *Glymnocladus* (Fig. 3). The distribution of Aze across three distinct orders supports a convergent evolutionary hypothesis for the independent emergence of Aze biosynthesis and future genomics and biochemical work could support this hypothesis.

## DISCUSSION

In this study we built phylochemical maps with NPAAs, which confirm and extend their known distribution across plants. Our analyses highlight the prevalence of NPAA production within legumes and additional orders where they are found. This approach can shed light on the evolutionary history of NPAA biosynthetic pathways and pinpoint species for additional biochemical and genomic studies.

### Increasing input data for phylochemical mapping

We chose 8 NPAAs as a starting point, mostly because of their common occurrence in the literature. However, of the ∼400,000 vascular plants (Christenhusz and Byng, 2016) only a small fraction have any literature evidence about the occurrence of NPAAs that were the focus of this study. To increase the utility of phylochemical mapping, two major improvements are needed, both involve increasing the amount of input data. Approaches that combine large language models and machine learning algorithms could be developed that not only search but assign metabolite-species associations with high accuracy and at large scales (Busta et al., 2024). This would enable identification of more NPAAs that are found across plants and provide a richer data set for phylochemical mapping. As a complimentary approach, metabolomics methods could be improved for larger scale detection of many NPAAs in a single analysis, similar to large scale screening of metabolites across a diverse genus (Ernst et al., 2019). Many of the methods used to detect NPAAs have been optimized for a single metabolite, thus most papers report association of one species one metabolite or a few species to the same metabolite. Methods should be developed so that NPAAs can be identified and quantified at scale, like methods used to quantify the 20 proteogenic amino acids (Thomas et al., 2024). Additionally NPAA-focused metabolomics approaches could provide true absence information of certain NPAAs, which addresses a limitation of literature-based phylochemical mapping. Simultaneous detection of many NPAAs from the same plant tissue will provide a holistic understanding of the types and distribution of NPAAs across plants.

### Do most plants not produce NPAAs?

Our focused analysis on a few NPAAs suggests that most orders do not accumulate NPAAs (Fig. 1). While this may be the case, we have mapped 8 of the more than 200 distinct NPAA structures identified in plants (Huang et al., 2011; Vranova et al., 2011). Before making conclusions about the restriction of NPAAs to certain species and orders a more global approach should be conducted. However, even from our focused analysis of NPAAs, it appears that some orders are more represented. For example, the Fabales, Malpighiales, Caryophyllales, and Poales all have multiple occurrences of structurally distinct NPAAs (Fig. 1). Additionally, legumes are well known for accumulation of structurally diverse NPAAs (Bell et al., 2008). This may suggest that these orders are predisposed to produce NPAAs, or perhaps NPAAs provide a selective advantage that is particularly important to these lineages. Mapping of more NPAAs into a phylogenetic context and in combination with genomic and biochemical studies could provide insight into the hypothesis that some lineages are predisposed for NPAA biosynthesis.

### Convergent evolution of some NPAA biosynthetic pathways

Convergent evolution is a common theme in plant specialized metabolism (Pichersky and Lewinsohn, 2011) with diverse metabolites arising independently in distinct lineages, including caffeine, betalains, and pyrrolizidine alkaloids (Reimann et al., 2004; Huang et al., 2016; Sheehan et al., 2020). Some NPAAs, such as Aze, are found in a few unrelated orders (Figs. 1, 2, & 3) and this distribution is highly suggestive of Aze biosynthesis independently emerging multiple times. Additionally, genera level phylogenies provide more resolution to when these pathways may have emerged (Fig. 3, green dots), and some instances into lineages that seem to have lost these pathways (Fig. 3 red dots). The lineages that have potentially lost Aze biosynthesis are equally interesting as those that have gained Aze biosynthesis and could provide insight into how and why metabolic pathways are lost. Metabolite distribution alone, however, only provides an indication of convergent evolution. Continued improvements and representation in the plant tree of life is crucial for accurate phylochemical mapping and interpretation (Zuntini et al., 2024).

### Phylochemical mapping to identify lineages for future investigations

Placement of metabolites into a phylogenetic context provides an indication of how pathways emerged and comparing biosynthetic pathways and the genes in distinct species may provide support for hypotheses about pathway evolution and loss events. Phylochemical mapping is a great first step, because it (1) leverages existing chemical data in the literature and (2) can be supplemented with chemical analysis with relative ease. Phylochemical mapping enables identification of plant lineages for follow-up experiments, which can be more challenging to perform at scale. Metabolite distributions together with complementary techniques such as genomics and biochemistry can provide a full understanding of the evolution of metabolic pathways.

As an example of using phylochemical mapping to identify species for future studies, here we identified an interesting distribution of djenkolic acid and mimosine (Fig. 2). Mimosine is narrowly distributed in two genera, *Mimosa* and *Leucaena*, whereas djenkolic acid is not reported in these genera, but all the genera surrounding (Fig. 2). Thus, mimosine-producing plants are embedded within the plants that produce djenkolic acid and the distribution of these two NPAAs appear to be mutually exclusive. It is possible that lack of data is partially responsible for the unique distribution of djenkolic acid and mimosine, however there could be metabolic changes in these species that enable one NPAA to be produced and not the other. As a first follow-up step, additional plants could be screened for presence of djenkolic acid and mimosine. Then, comparative analyses using genomics and biochemistry of the legume lineages that produce djenkolic acid and mimosine could provide insight into how djenkolic acid and mimosine evolved in these legume lineages and if these compounds are mutually exclusive.

Phylochemical mapping approaches can be applied to metabolites distributed across organisms in any taxonomic grouping. Given the high-quality genomes of the tree of life, this approach can provide insight into how and why lineages produce specific metabolites. Continued improvements in metabolomics, genomics, systematics, and high-throughput literature scanning approaches will enhance the power of phylochemical mapping and strengthen the conclusions that can be drawn from modern-day chemotaxonomy.

## AUTHOR CONTRIBUTIONS

MG performed wet lab and computational experiments, analyzed data, created figures and reviewed and revised the text. WTS performed wet lab and computational experiments, analyzed data, created figures and reviewed and revised the text. AO performed literature searches, compiled data, and edited the text. LB conceptualized the project, analyzed data, created figures and edited the text. CAS conceptualized the project, analyzed data, created figures, wrote and edited the text.

## ACKNOWLEDGEMENTS

We thank Monica Carlsen and Meghan Forde at the Missouri Botanical Garden for help in collecting plant tissue. We acknowledge the following botanical gardens that have provided the Missouri Botanical Garden with plant material that was used in our study: Jardí Botànic de Barcelona, Botanical Garden of the Institute of Ecology and Botany, Vácrátót, and Jerusalem Botanical Gardens. We thank Pete Millier and Leanne Tippett Mosby at the University of Missouri for help collecting plant material from our campus botanical garden. We thank George Frees for help with plant tissue collection. We thank the USDA for providing diverse germplasm. CAS acknowledges financial support for this project from a University of Missouri College of Agriculture, Food and Natural Resources Joy of Discovery Grant. L.B. was financially supported by and gratefully acknowledges the Swenson College of Science and Engineering at the University of Minnesota Duluth.

## DATA AVAILIBILITY STATEMENT

All the relevant data and code are available in the supplementary materials or on our GitHub page https://github.com/thebustalab/npaa_distribution.

## TITLES FOR SUPPORTING INFORMATION

**Supplementary Figure 1.**
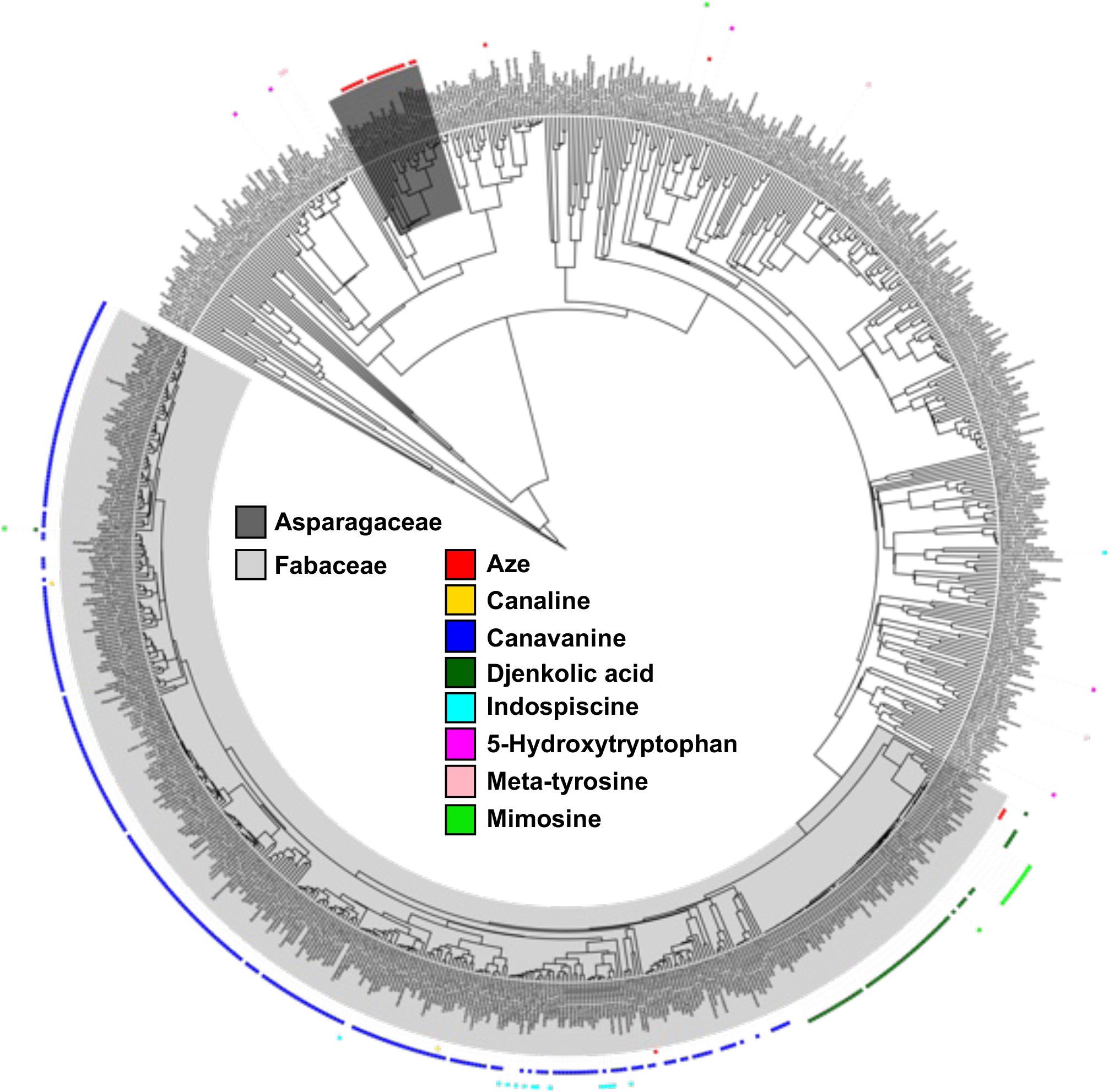
Species level distribution of NPAAs across plants with species labels. Same figure as is shown in Fig. 2, but with genus species names for each tip of the tree shown.

## REFERENCES

Alseekh, S., and A. R. Fernie. 2018. Metabolomics 20 years on: what have we learned and what hurdles remain? The Plant Journal: For Cell and Molecular Biology 94: 933–942.

Alston, R. E., R. E. Alston, and B. L. Turner. 1963. Biochemical systematics. Prentice-Hall, Englewood Cliffs, N.J.

Bate-Smith, E. C. 1962. The phenolic constituents of plants and their taxonomic significance. I. Dicotyledons. Botanical Journal of the Linnean Society 58: 95–173.

Beaudoin, G. A. W., and P. J. Facchini. 2014. Benzylisoquinoline alkaloid biosynthesis in opium poppy. Planta 240: 19–32.

Bell, E. A. 2003. Nonprotein amino acids of plants: significance in medicine, nutrition, and agriculture. Journal of Agricultural and Food Chemistry 51: 2854–2865.

Bell, E. A. 1976. ‘Uncommon’ amino acids in plants. FEBS letters 64: 29–35.

Bell, E. A., A. A. Watson, and R. J. Nash. 2008. Non-protein amino acids: a review of the biosynthesis and taxonomic significance. Natural Product Communications 3: 1934578X0800300.

Bertin, C., L. A. Weston, T. Huang, G. Jander, T. Owens, J. Meinwald, and F. C. Schroeder. 2007. Grass roots chemistry: meta-Tyrosine, an herbicidal nonprotein amino acid. Proceedings of the National Academy of Sciences 104: 16964–16969.

Busta, L., D. Hall, B. Johnson, M. Schaut, C. M. Hanson, A. Gupta, M. Gundrum, et al. 2024. Mapping of specialized metabolite terms onto a plant phylogeny using text mining and large language models. The Plant Journal: For Cell and Molecular Biology.

Christenhusz, M. J. M., and J. W. Byng. 2016. The number of known plants species in the world and its annual increase. Phytotaxa 261: 201–217.

Dixon, R. A., and D. Strack. 2003. Phytochemistry meets genome analysis, and beyond. Phytochemistry 62: 815–816.

Endara, M.-J., D. L. Forrister, and P. D. Coley. 2023. The Evolutionary Ecology of Plant Chemical Defenses: From Molecules to Communities. Annual Review of Ecology, Evolution, and Systematics 54: 107–127.

Erb, M., and D. J. Kliebenstein. 2020. Plant secondary metabolites as defenses, regulators, and primary metabolites: The blurred functional trichotomy. Plant Physiology 184: 39–52.

Ernst, M., L.-F. Nothias, J. J. J. van der Hooft, R. R. Silva, C. H. Saslis-Lagoudakis, O. M. Grace, K. Martinez-Swatson, et al. 2019. Assessing Specialized Metabolite Diversity in the Cosmopolitan Plant Genus Euphorbia L. Frontiers in Plant Science 10: 846.

Fowden, L. 1955. Azetidine-2-carboxylic acid: a new constituent of plants. Nature 176: 347–348.

Fowden, L., and F. C. Steward. 1957. Nitrogenous compounds and nitrogen metabolism in the Liliaceae: I. The occurrence of soluble nitrogenous compounds. Annals of Botany 21: 53–67.

Gibbs, R. D. 1974. Chemotaxonomy of Flowering Plants: Four Volumes. McGill-Queen’s University Press.

Grobbelaar, N., J. K. Pollard, and F. C. Steward. 1955. New soluble nitrogen compounds (amino- and imino-acids and amides) in plants. Nature 175: 703–708.

Huang, R., A. J. O’Donnell, J. J. Barboline, and T. J. Barkman. 2016. Convergent evolution of caffeine in plants by co-option of exapted ancestral enzymes. Proceedings of the National Academy of Sciences of the United States of America 113: 10613–10618.

Huang, T., G. Jander, and M. de Vos. 2011. Non-protein amino acids in plant defense against insect herbivores: representative cases and opportunities for further functional analysis. Phytochemistry 72: 1531–1537.

Leebens-Mack, J. H., M. S. Barker, E. J. Carpenter, M. K. Deyholos, M. A. Gitzendanner, S. W. Graham, I. Grosse, et al. 2019. One thousand plant transcriptomes and the phylogenomics of green plants. Nature 574: 679–685.

Letunic, I., and P. Bork. 2024. Interactive Tree of Life (iTOL) v6: recent updates to the phylogenetic tree display and annotation tool. Nucleic Acids Research 52: W78–W82.

Li, F.-S., and J.-K. Weng. 2017. Demystifying traditional herbal medicine with modern approach. Nature Plants 3: 17109.

McChesney, J. D., S. K. Venkataraman, and J. T. Henri. 2007. Plant natural products: back to the future or into extinction? Phytochemistry 68: 2015–2022.

Moghe, G. D., and R. L. Last. 2015. Something old, something new: conserved enzymes and the evolution of novelty in plant specialized metabolism. Plant Physiology 169: 1512–1523.

Newman, D. J., and G. M. Cragg. 2016. Natural Products as Sources of New Drugs from 1981 to 2014. Journal of Natural Products 79: 629–661.

Pichersky, E., and E. Lewinsohn. 2011. Convergent evolution in plant specialized metabolism. Annual Review of Plant Biology 62: 549–566.

Qian, H., and Y. Jin. 2016. An updated megaphylogeny of plants, a tool for generating plant phylogenies and an analysis of phylogenetic community structure. Journal of Plant Ecology 9: 233–239.

Rai, A., K. Saito, and M. Yamazaki. 2017. Integrated omics analysis of specialized metabolism in medicinal plants. The Plant Journal: For Cell and Molecular Biology 90: 764–787.

Reimann, A., N. Nurhayati, A. Backenköhler, and D. Ober. 2004. Repeated evolution of the pyrrolizidine alkaloid-mediated defense system in separate angiosperm lineages. The Plant Cell 16: 2772–2784.

Reynolds, T. 2007. The evolution of chemosystematics. Phytochemistry 68: 2887–2895.

Rodriguez-Mias, R. A., K. N. Hess, B. Y. Ruiz, I. R. Smith, A. S. Barente, S. M. Zimmerman, Y. Y. Lu, et al. 2022. Proteome-wide identification of amino acid substitutions deleterious for protein function. 2022.04.06.487405.

Schenck, C. A., and R. L. Last. 2020. Location, location! cellular relocalization primes specialized metabolic diversification. The FEBS Journal 287: 1359–1368.

Sheehan, H., T. Feng, N. Walker-Hale, S. Lopez-Nieves, B. Pucker, R. Guo, W. C. Yim, et al. 2020. Evolution of L-DOPA 4,5-dioxygenase activity allows for recurrent specialisation to betalain pigmentation in Caryophyllales. The New Phytologist 227: 914–929.

Sung, M.-L., and L. Fowden. 1969. Azetidine-2-carboxylic acid from the legume Delonix regia. Phytochemistry 8: 2095–2096.

Thives Santos, W., V. Dwivedi, M. Miederhoff, K. V. Hoek, H. Duong, R. Angelovici, and C. Schenck. 2024. Mode of action of the toxic proline mimic azetidine 2-carboxylic acid in plants. bioRxiv.

Thomas, S. K., K. V. Hoek, T. Ogoti, H. Duong, R. Angelovici, J. C. Pires, D. Mendoza-Cozatl, et al. 2024. Halophytes and heavy metals: A multi-omics approach to understand the role of gene and genome duplication in the abiotic stress tolerance of Cakile maritima. American Journal of Botany: e16310.

Vendemiatti, E., L. Nowack, L. E. P. Peres, V. A. Benedito, and C. A. Schenck. 2024. Sticky business: the intricacies of acylsugar biosynthesis in the Solanaceae. Phytochemistry Reviews.

Vranova, V., K. Rejsek, K. R. Skene, and P. Formanek. 2011. Non-protein amino acids: plant, soil and ecosystem interactions. Plant and Soil 342: 31–48.

Weng, J.-K., J. H. Lynch, J. O. Matos, and N. Dudareva. 2021. Adaptive mechanisms of plant specialized metabolism connecting chemistry to function. Nature Chemical Biology 17: 1037–1045.

Wickham, H. 2009. ggplot2: Elegant Graphics for Data Analysis. Springer, New York, NY.

Yu, G., D. K. Smith, H. Zhu, Y. Guan, and T. T.-Y. Lam. 2017. ggtree: an r package for visualization and annotation of phylogenetic trees with their covariates and other associated data. Methods in Ecology and Evolution 8: 28–36.

Zer, H., H. Mizrahi, N. Malchenko, T. Avin-Wittenberg, L. Klipcan, and O. Ostersetzer-Biran. 2020. The phytotoxicity of meta-tyrosine is associated with altered phenylalanine metabolism and misincorporation of this non-proteinogenic Phe-analog to the plant’s proteome. Frontiers in Plant Science 11: 140.

Zuntini, A. R., T. Carruthers, O. Maurin, P. C. Bailey, K. Leempoel, G. E. Brewer, N. Epitawalage, et al. 2024. Phylogenomics and the rise of the angiosperms. Nature 629: 843–850.

